# Incorporating calibrated functional assay data into the *BRCA1* Ex-UV database

**DOI:** 10.1101/079418

**Authors:** Bryony A. Thompson, Russell Bell, Bryan E. Welm, John Burn, Sean V. Tavtigian

## Abstract

Driven by massively parallel sequencing and allied technologies, the scale of genetic predisposition testing is on a dramatic uptrend. While many patients are found to carry clinically actionable pathogenic sequence variants, testing also reveals enormous numbers of Unclassified Variants (UV), or Variants of Uncertain Significance (VUS), most of which are rare missense substitutions. Following IARC variant classification guidelines, quantitative methods have been developed to integrate multiple data types for clinical UV evaluation in *BRCA1/2*; results from these analyses are recorded in the BRCA gene Ex-UV database (hci-exlovd.hci.utah.edu). In variant classification, the rate-limiting step is often accumulation of patient observational data. Recently, functional assays evaluating *BRCA1* RING domain and C-terminal substitutions have been calibrated, enabling variant classification through a two-component combination of sequence analysis-based predictions with functional assay results. This two-component classification was embedded in a decision tree with safeguards to avoid misclassification. For the two-component analysis, sensitivity is 87.5%, specificity is 100%, and the error rate 0.0%. Classification of a UV as likely pathogenic or likely neutral does not require certainty; the probabilistic definitions of the categories imply an error rate. Combining sequence analysis with functional assay data in two-component analysis added 146 *BRCA1* variants to the Ex-UV database.

## Introduction

Inactivating mutations in *BRCA1* (MIM #113705) are associated with an increased risk of breast and ovarian cancer. While many patients who undergo clinical mutation testing are found to carry clinically actionable pathogenic sequence variants, testing also reveals large numbers of Unclassified Variants (UV), or Variants of Uncertain Significance (VUS), most of which are rare missense substitutions. All known pathogenic non-spliceogenic *BRCA1* missense substitutions are found in the N-terminal RING domain and C-terminal BRCT domains (Easton et al. 2007; Lindor et al. 2012; Vallée et al. 2012). The RING domain binds BARD1, enabling rapid recruitment to sites of DNA damage and generating an E3 ubiquitin ligase (Li and Yu 2013; Densham et al. 2016), and the BRCT domains bind a number of proteins involved in the DNA damage response (reviewed in Mermershtain and Glover 2013). Many functional assays have been developed to test the impact of missense substitutions on these functionally important domains, and other regions of the BRCA1 protein.

The BRCA gene Ex-UV database (hci-exlovd.hci.utah.edu) contains *BRCA1* and *BRCA2* UVs clinically classified or reclassified using a quantitative Integrated Evaluation (QIE, also known as multifactorial likelihood analysis) (Vallée et al. 2012). The classifications are based on calibrated sequence analysis to generate a prior probability of pathogenicity (Prior P), patient observational data to generate odds in favor of pathogenicity, and Bayes rule to combine these data into a posterior probability of pathogenicity (Posterior P). For purposes of clinical translation and patient counselling/management, the Posterior P is translated through the IARC 5-category classifier (Plon et al. 2008). Currently the database includes fields for sequence analysis-based Prior Ps for missense analysis (Tavtigian et al. 2008), and predicted effects on splicing (Vallée et al. 2016). It also includes odds in favor of pathogenicity in the form of likelihood ratios (LR) for co-segregation with cancer phenotype (Thompson et al. 2003), strength of personal and family cancer history (Easton et al. 2007), tumor pathology (Chenevix-Trench et al. 2006; Spurdle et al. 2008, 2014), and co-occurrence with known pathogenic variants (Goldgar et al. 2004); the product of these four LRs is the observational LR in favor of pathogenicity (Obs LR).

The rate-limiting step in UV classification has been accumulation of sufficient Obs LR from patient observational data to modify the Prior P into a Posterior P >0.95 (likely pathogenic) or <0.05 (likely not pathogenic/likely neutral). Functional assay estimates of damage to protein function can be used as a proxy for pathogenicity if the assay outputs are empirically calibrated (either against known pathogenic and known neutral variants or against patient observational data) to read out a variable that is more directly related to pathogenicity e.g., a functional likelihood ratio in favor of pathogenicity (Functional LR). Once such a calibration is in place and the functional assay(s) are considered sufficiently accurate for use in clinical variant classification, the assay(s) can become directly useful for medical genetics related evaluation of missense substitutions and other UVs. In principle, the QIE enables classification through a two-component combination of sequence-based *in silico* analyses and a calibrated functional assay.

In the companion article (Paquette et al., submitted and bioRxiv 092619; doi: http://dx.doi.org/10.1101/092619) we describe the calibration of a mammalian two-hybrid assay testing BARD1 binding capabilities of BRCA1 RING domain substitutions. In addition, Woods *et al*. (2016) recently described the calibration of mammalian transcriptional activation (TA) assays for evaluation of missense substitutions in the BRCA1 C-terminal region. Here we report an update of the *BRCA1* Ex-UV database to incorporate a Functional LR based on these calibrated functional assays into the Integrated Evaluation of *BRCA1* sequence variants. This has led to the addition of 152 variants to the database.

## Methods

### MIM # and accession #

*BRCA1* is MIM# 113705, and exonic variant coding used here is based on NM_007294.3.

### Database updates

Three new fields have been added to the LOVD *BRCA1* Ex-UV database (hciexlovd.hci.utah.edu, under the gene database BRCA1fx): Functional LR, Functional Assay Reference, and Clinical Severity. The Clinical Severity field will be used to differentiate pathogenic variants (i.e. confers medically actionable increased risk) associated with different levels of risk. The values used to populate Clinical Severity are high-risk, moderate-risk, and not applicable. The Clinical Severity category is assigned to a variant based on the estimated cumulative risks to age 80: high-risk >32%; moderate-risk 18-32% (Easton et al. 2015). Variants assigned to Class 5 or Class 4 by QIE are assumed to be high-risk unless patient observational data in the form of co-segregation with the cancer phenotype, strength of personal and family cancer history, or a case-control odds ratio are indicative of moderate-risk. At this time, p.R1699Q is the only proven moderate-risk *BRCA1* variant (Spurdle et al. 2012). Clinical Severity for variants assigned to Class 3, Class 2, or Class 1 by QIE are marked not applicable. The other functional assay-related LOVD fields are populated based on the functional assay data available in the ‘datasets’ section described below.

### Datasets

Only missense substitutions that have been observed in cases were added to the LOVD database, due to assumed contribution of a minimal observational datum (personal history of cancer) in the derivation of the sequence analysis Prior P and Functional LR. For the BRCA1 RING domain (aa 2–103), there were 10 natural (i.e. originally identified in cancer cases) already classified missense substitutions recorded in the Ex-UV database, and an additional UV (p.P34S). Because only two variants were neutral, the assay calibration described in the companion research article (Paquette et al., submitted and bioRxiv 092619; doi: http://dx.doi.org/10.1101/092619) included three human Class 3 substitutions with Obs LR <0.50, plus four cross-species multiple sequence alignment neutral missense substitutions *(viz*, the alternate amino acid is present in a non-human primate and within the range of variation at it’s position in a mammals-only BRCA1 protein multiple sequence alignment even if the specific primate with that residue is excluded from the alignment). To avoid circularities, the variants used in assay calibration in Paquette *et al*. (submitted and bioRxiv 092619; doi: http://dx.doi.org/10.1101/092619) are not included here. There are 242 ‘natural’ *BRCA1* missense substitutions between amino acids 1,315-1,863 evaluated in Woods *et al*. (2016), including 36 variants that were used as internal standards in calibration of the transactivation assay. These were in Lindor *et al*. (2012) as IARC Class 1, Not Pathogenic (n=19); Class 4, Likely Pathogenic (n=1); and Class 5, Pathogenic (n=14). One of the ‘neutral’ variants (c.4484G>T p.R1495M) is actually a spliceogenic pathogenic variant (Colombo et al. 2013), and the moderate-risk variant p.R1699Q was assigned to Class 5. A further two Class 1 variants were classified by the Evidence-based Network for the Interpretation of Germline Mutant Alleles (ENIGMA) Consortium (data not shown). As with the RING domain variants these were not added to the BRCA1fx database to avoid circularities. The remaining 206 variants include 12 in the coiled-coil domain (aa 1,392-1,424); 117 over the BRCT1, linker, and BRCT2 domains (aa 1,650-1,859); and 73 in the interval between the coiled-coil and BRCT1. The missense analysis Prior P and splicing Prior P for each variant are taken from the HCI Breast Cancer Genes Prior Probabilities database (priors.hci.utah.edu/PRIORS) (Vallée et al. 2016).

### Derivation of Functional LR

The derivation of the functional Odds Path for the BRCA1 RING domain missense substitutions is described in the companion article (Paquette et al submitted and bioRxiv 092619; doi: http://dx.doi.org/10.1101/092619). For the missense substitutions in Woods *et al*. (2016) the supplementary materials provided the data used to derive the Functional LR. They used a Bayesian hierarchical model (called VarCall) that takes into account experimental heterogeneity in the *in cellulo* functional assay to estimate log_e_ odds in favor of pathogenicity (L_e_LR) for each variant tested (Iversen et al. 2011; Woods et al. 2016).

Unlike the mammalian two-hybrid assay described in the companion article (Paquette et al submitted and bioRxiv 092619; doi: http://dx.doi.org/10.1101/092619), the TA assay does not directly test the underlying mechanism causing pathogenicity of a missense substitution in the C-terminus. For example, it is unable to discriminate the moderate-risk pathogenic variant *BRCA1* p.R1699Q from high-risk pathogenic variant p.R1699W. Thus, we wanted to introduce a degree of uncertainty to the calculation of the Functional LR, through constraining the L_e_LR to act as a safe-guard against misclassification based on evaluation by two-component QIE. Previously classified variants on the Ex-UV database (n=38, Supp. Table S1) were used to define the zone of uncertainty, which informed the value used to constrain the log_e_ odds estimated by VarCall. The constrained Functional LR was then calculated by exponentiation of the constrained L_e_LR.

### Statistical analyses

Sensitivity was calculated as (# true pathogenic variants detected by a classifier/total # true pathogenic variants). Specificity was calculated as (# true neutral variants detected by a classifier/total # true neutral variants). Error rate was calculated as [(# false pathogenic variants + # false neutral variants detected by a classifier)/(total # true pathogenic variants + # true neutral variants)]. 95% confidence intervals on these proportions were estimated using the binomial confidence interval calculator in STATA 13.1 (StataCorp).

We explored how sensitive two-component QIE would be to misspecification of either the missense analysis Prior Ps or the constrained Functional LRs. For this analysis, we assumed that the total number of misclassified variants could be estimated from their Posterior Ps, if these were accurate. For groups of pathogenic variants, the estimated number or errors would be given by equation 1, and for groups of neutral variants by equation 2:

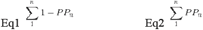

Where ***n*** is an index number for each variant in the group and ***PP*** is the corresponding Posterior P. The misclassification rate would simply be the estimated number of misclassified variants divided by ***n***, with the understanding that a rate >0.05 would violate the IARC guideline for the likely neutral or likely pathogenic categories.

## Results and Discussion

### Embedding quantitative integrated evaluation, with inclusion of calibrated functional assays, in a decision tree

Sequence analysis-based Prior Ps are available for all missense substitutions in *BRCA1* that can be reached by a single nucleotide substitution. Any of these substitutions can be subjected to QIE if one or more of four types of patient observational data are available: personal and family cancer history, co-segregation with cancer phenotype, co-occurrence between UVs and clearly pathogenic variants, and/or tumor immunohistochemistry & grade (Lindor et al. 2012; Vallée et al. 2012). Here, we add calibrated function assays to the mix. To guide the analysis of individual substitutions while providing some safeguards against misclassification, we embed the QIE in a decision tree (Fig. 1). Logic behind key nodes of the tree is explained in the next few paragraphs. The discussion proceeds from the assumption that valid functional assay data exists (node 1).

**Figure 1.**
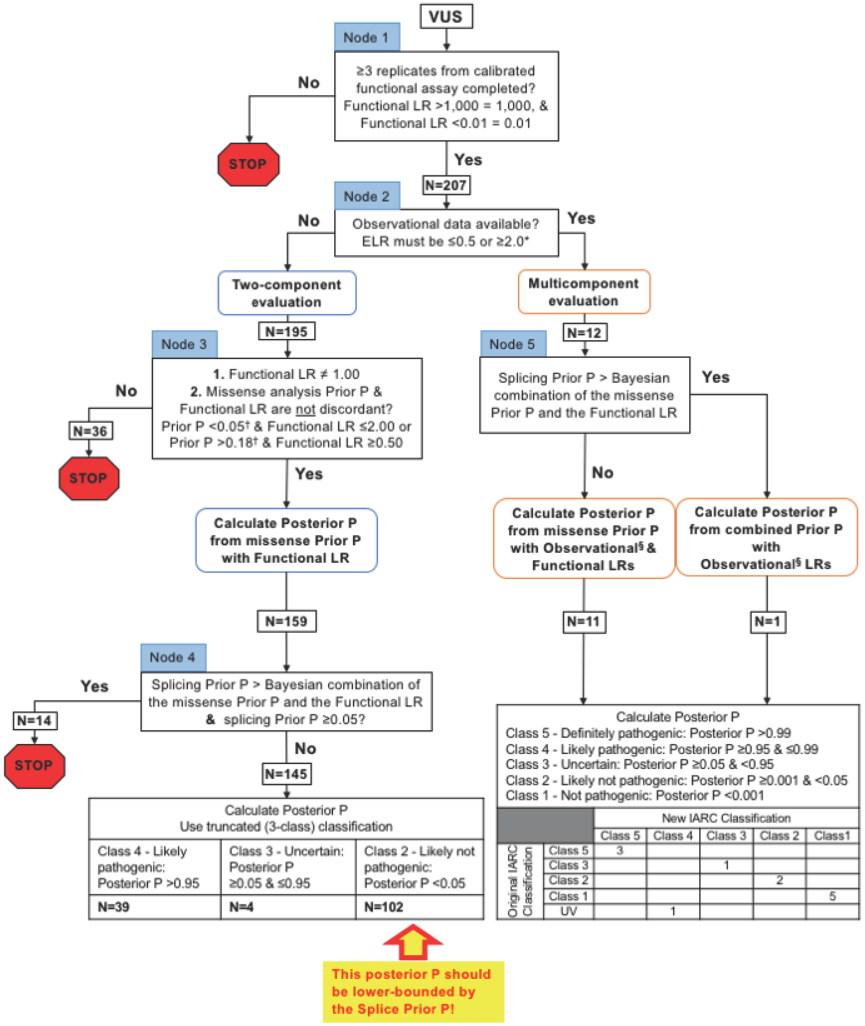
Functional assay decision tree. N-number of *BRCA1* missense substitutions. P-probability of pathogenicity. *There must be enough subjects/observations that if the variant were pathogenic, then the expected LR (ELR) would be >2, or if the variant were neutral, the ELR would be <0.5. ^†^Two-fold either side of 0.1, the overall sequence-based analysis prior for *BRCA1/2* (see main text for explanation). ^§^Observational LRs currently include: segregation; tumor pathology; summary family history; and co-occurrence.

Inclusion of patient observational data (node 2). Recently, Vallee *et al*. asked the philosophical question of how much observational data are required in order to perform a *bona fide* integrated evaluation, and concluded that Obs LR of ≤0.5 or ≥2.0 are reasonable inner boundaries for the magnitude of observational data required (Vallée et al. 2016). Noting that some variants, especially moderate-risk variants such as *BRCA1* p.R1699Q can be resistant to classification because the observational data can be conflicting, a more refined criterion would be that a multicomponent QIE with observational data requires either that the Obs LR criterion of ≤0.5 or ≥2.0 is met, or else at least enough observational data to expect an Obs LR ≥2.0 if the variant were pathogenic. As examples of the expected Obs LR, tumor estrogen receptor (ER) status is an extremely effective data type because ~75% of breast tumors from *BRCA1* mutation carriers are ER-, and estrogen receptor negativity provides an Obs LR of 2.6 in favor of pathogenicity (so long as the case was not ascertained because of their tumor's immunohistochemical profile) (Spurdle et al. 2014). Therefore, data from just one tumor is enough to meet the expected Obs LR criterion. At the other extreme, analysis of co-observation between UVs and clearly pathogenic variants is relatively ineffective for classification as pathogenic because a specific *BRCA1* UV has to be observed about 17 times in the absence of a pathogenic variants in order to generate an Obs LR of 2.0 in favor of pathogenicity (Goldgar et al. 2004; Tavtigian et al. 2006). If the Obs/expected Obs LR criterion is met, multicomponent QIE is applied; otherwise, analysis moves to node 3 and a two-component evaluation as described in the next paragraph.

Currently there are two mammalian functional assays evaluating the impact of *BRCA1* missense substitutions that have been calibrated. These are a mammalian two-hybrid assay testing the RING domain (Paquette et al submitted and bioRxiv 092619; doi: http://dx.doi.org/10.1101/092619), and a TA assay used to test many reported missense substitutions in the BRCA1 C-terminal region (Iversen et al. 2011; Woods et al. 2016). At the first step of two-component classification (node 3), two simple safeguards against error could be that (1) the Functional LR cannot equal 1.00, and (2) if the missense analysis Prior P and the Functional LR are discordant, then classification is not permitted. The first point ensures that the Prior P is not directly converted to a Posterior P. Accordingly, for the second point, consider a variant where the missense analysis Prior P is 0.49 and the Functional LR is 1.04. One might say that these data are discordant because they are on opposite sides of neutrality, but both results indicate uncertainty. So, it would be perfectly reasonable to combine these into a Posterior P, which would be 0.50. A more interesting situation would be if the missense analysis Prior P gave 2-fold evidence against pathogenicity (naively, Prior P = 0.33) and functional assay gave 2fold evidence in favor (Functional LR = 2.0). Here, we think that it’s fair to say that they “disagree”. So, we propose that if the missense analysis Prior P and Functional LR are on opposite sides of a factor-of-2 window around neutrality, we will NOT combine them via a two-component QIE, and the missense substitution remains Unclassified. For *BRCA1*, a small complication is that the background Prior P for an individual rare missense substitution is approximately 0.1, not 0.5 (Abkevich et al. 2004; Goldgar et al. 2004). Thus, the definitions for non-discordant combinations are missense analysis Prior P <0.05 and Functional LR ≤2.00, or Prior P >0.18 and Functional LR ≥0.50 (Fig. 1).

In general, there are two ways for a missense substitution to be pathogenic: the missense substitution can damage a key function of the protein, or the underlying nucleotide substitution can alter mRNA splicing in a way that results in either a non-productive transcript or damage to a key function of the protein. The missense analysis Prior P and Functional LR only address damage to a key function of the protein. Accordingly, consider a missense substitution where the Bayesian combination of the missense analysis Prior P and Functional LR naively results in a probability of pathogenicity <0.05, but the mRNA splicing analysis Prior P (Vallée et al. 2016) is >0.05 (Fig. 1, node 4). Since the functional assays evaluated here do not assess spliceogenicity, the overall probability of pathogenicity remains >0.05 and is given by the splicing Prior P. In this case, the missense substitution remains Unclassified. Note that this issue is explicitly asymmetric. If the Bayesian combination of the missense analysis Prior P and Functional LR results in a probability of pathogenicity >0.95, but the splicing Prior P <0.05, the substitution has Post P >0.95 and can be considered IARC Class 4, Likely Pathogenic. For missense substitutions that pass node 4, the Post P is used to place the substitution on a truncated 3 Class version of the IARC classifier.

> Class 4-Likely pathogenic: Posterior P >0.95
>
> Class 3 - Uncertain: Posterior P ≥0.05 & ≤0.95
>
> Class 2 - Likely not pathogenic: Posterior P <0.05

Moving back to node 2, if the observational/expected Obs LR criterion is met, multicomponent QIE can be applied and the analysis moves to node 5. Here, the question is much the same as at node 4: which is higher, the Bayesian combination of the missense analysis Prior P and Functional LR, or the splicing Prior P? If the splicing Prior P is greater, suggesting that there is a high probability the variant is spliceogenic, the Obs LR from patient data only is applied to this probability to calculate the missense substitution’s Post P. Otherwise, the missense analysis Prior P is applied to the Obs LR derived from patient data and Functional LR. The Post P is then used to place the substitution in the standard 5 Class IARC classifier (Plon et al. 2008).

### Safeguards against UV misclassification built into the decision tree and quantitative integrated evaluation with functional assay likelihood ratios

As integrated within the decision tree of Figure 1, QIE with Functional LRs includes five specific safeguards designed to reduce the risk of UV misclassification. These safeguards will also result in some UVs that would have been correctly classified instead resting as Unclassified pending accumulation of further data.

1. The Functional LR for variants with severe loss of function is capped at 1,000 (L_e_LR = 6.9), and the Functional LR for a variant with full function is capped at 0.01 (L_e_LR = -4.6). The ceiling and floor prevent stand-alone reversal of classification by the Functional LR in a multi-component QIE including the combined prior and observational data. For example, a key domain Align-GVGD C65 missense substitution has a missense Prior P of 0.81. If such a variant had a minimally concordant Obs LR of 2.0, it would have a Posterior P of 0.895–trending towards but not quite reaching Likely Pathogenic. But a Functional LR of ≤0.0061 (strongly indicative of neutrality) would result in a new Posterior P of <0.050, reversing the combination of Prior P and observational data to a classification of Likely Neutral. A Functional LR cap at 0.01 prevents this stand-alone reversal. Similarly, a key domain Align-GVGD C0 missense substitution has a missense Prior P of 0.03. If such a variant had an Obs LR of 0.5, it would be classified as Class 2 Likely Not Pathogenic with a Posterior P of 0.0152. But a Functional LR of 1,230 would drive the Posterior P above 0.950 and reverse the classification to Likely Pathogenic. The ceiling and floor are somewhat *ad hoc*, tuned to the actual missense Priors Ps and minimal Obs LR data inclusion rules already embedded in the QIE (Tavtigian et al. 2008; Vallée et al. 2016). Nonetheless, they also correspond exactly to the stand-alone classification criteria from the original framework for quantitative classification of *BRCA1/2* UVs (Goldgar et al. 2004).
2. Data from the classified variants used to train VarCall (n=28, Supp. Table S1) Iversen et al. (2011) or Woods et al. (2016) presented a notable lack of variants with L_e_LR between ^-^2 and +2 (Fig. 2A). This means that the *%* wildtype activity corresponding to the tipping point where the TA assay result switches from evidence against pathogenicity to evidence for pathogenicity was interpolated from variants with activities well above and below that point, rather than being informed by variants with activities near that point. To accommodate uncertainty in the tipping point, we inserted an interval with constrained L_e_LR=0 into the Woods et al. (2016) L_e_LR between its native scores of ^-^2 and +2. Similar to the functional floor and ceiling described in point 1 above, the offset of 2.0 is somewhat *ad hoc*, because unlike the BRCA1 RING domain assay (Paquette et al submitted and bioRxiv 092619; doi: http://dx.doi.org/10.1101/092619), we were not able to derive confidence intervals for the TA assay calibration. Figure 2B, corresponding to Table 1, shows the conversion from the reported LeLR to constrained LeLR for the missense substitutions tested in the C-terminus. After exponentiation of the constrained L_e_LR to derive the constrained Functional LR, the variants can enter the decision tree.
3. Unclassified Variants are remanded for further analysis if their missense Prior P and Functional LR are discordant. For the missense analysis Prior P, there are clear instances of substitutions with low Prior Ps that are nonetheless pathogenic because of missense dysfunction, and clear instances of substitutions with high missense Prior Ps that are clearly neutral or nearly so. On the functional side, *BRCA1* p.R1699Q has a more damaging effect in the TA assay than does p.R1699W. Yet evidence from personal and family cancer histories are consistent with p.R1699Q conferring a moderate-risk phenotype whereas p.R1699W is consistent with a high-risk phenotype (Easton et al. 2007; Spurdle et al. 2012). Since it is therefore clear that the Functional LR is not a perfect predictor of disease risk, the discordance rule is a prudent precaution, albeit with a somewhat arbitrary definition.
4. The *BRCA1* missense substitution p.L1407P presents a more interesting issue: misclassification of the variant because of a key domain specification issue for the missense analysis Prior P. The substitution falls in the BRCA1 coiled-coil domain (Fig. 3) that mediates interaction between BRCA1 and PALB2. It is predicted to be deleterious (align GVGD score = C65), and demonstrates an intermediate effect in functional assays (constrained Functional LR 1.62). Similarly, *BRCA1* p.M1411T is predicted to be deleterious, but falls into the grey zone around intermediate function in the TA assay (Fig 2B, constrained Functional LR 1.00). Both variants abrogate PALB2 binding to the coiled-coil domain, and there is evidence that damaging this interaction compromises repair of DNA double strand breaks by homologous recombination (Sy et al. 2009; Woods et al. 2016). Nonetheless, it is not currently known whether missense substitutions that disrupt the function of this coiled-coil domain are associated with an increased cancer risk, and missense substitutions in this domain are assigned missense Prior Ps of 0.02 (Easton et al. 2007; Tavtigian et al. 2008). Patient observational data could be used to answer the question, if data were available from several pedigrees that segregate these variants-which is exactly the rate-limiting problem for variant classification. Alternatively, specific mouse knock-ins of this class of missense substitution, e.g. Align-GVGD C65 substitutions that also show clear damage in a functional assay, could be used to estimate penetrance compared to known reference sequence and pathogenic controls (Shakya et al. 2011). The same issue arises for missense substitutions in the *BRCA2* (exon 2-3) PALB2 interaction domain and (exon 27) RAD51 disassembly domain (Xia et al. 2006; Davies and Pellegrini 2007).
5. Unclassified Variants are also remanded for further analysis if their splicing Prior P is ≥0.05 and their splicing Prior P is > the Bayesian combination of the missense analysis Prior P and the Functional LR. Potentially spliceogenic missense substitutions that could damage a splice donor, damage a splice acceptor, or create a *de novo* donor are caught by this safeguard, specifically reducing the probability of false negative classification through QIE.

**Figure 2.**
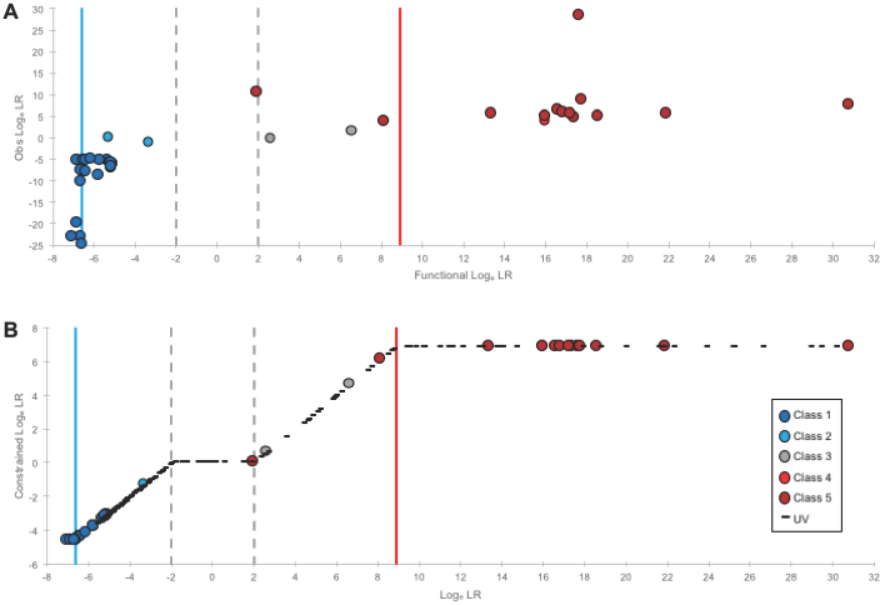
Estimation of constrained log_e_ likelihood ratios for safeguard against UV misclassification. A: Comparison of the reported functional assay log_e_ LR to log_e_ transformed Obs LR derived from patient observational data (Product LR in the Ex-UV database) for previously classified variants (n=38) used to define the zone of uncertainty of the transcriptional activation assay. B: Conversion to constrained log_e_ LR for BRCA1 C-terminus missense substitutions (n=233). The IARC classifications of the variants are defined in the legend. The dashed lines define the boundaries of the zone of uncertainty, which was used to define the magnitude at which the log_e_ LR (L_e_LR) would be constrained (± 2). The solid lines indicate the L_e_LR equivalent to the Functional LR ceiling for severe loss of function (constrained Functional LR = 1,000; constrained L_e_LR = 6.9; L_e_LR = 8.9), and floor for full function (constrained Functional LR = 0.01; constrained L_e_LR = -4.6; L_e_LR = -6.6).

**Figure 3.**
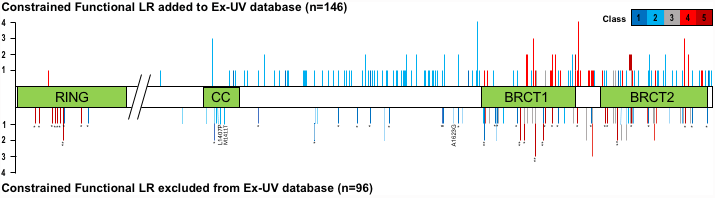
Map of *BRCA1* missense substitutions tested in functional assays. A schematic of the missense substitutions tested in the RING domain and the C-terminal of the BRCA1 protein color-coded by the 5-category classifier. Includes variants evaluated (n=146) using two-component and multicomponent quantitative integrated evaluation including the constrained Functional LR, as outlined in Figure 1. The remaining variants (n=96) include the missense substitutions used in calibration of the functional assays (n=46, color-coded as pathogenic or neutral), and substitutions that could not be classified using the constrained Functional LR (n=50, color-coded to what the assay results indicated based on two-component evaluation). The dashed lines are the variants where the splicing predictions overruled the results of two-component evaluation. Locations of domains: RING-aa 2-103; CC, coiled-coil-aa 1,3921,424; BRCT1 - aa 1,650-1,736; BRCT2 - aa 1,760-1,859. * - variants used in assay calibration.

### Performance on previously classified variants with patient observational data

To get some idea of the performance of classification based on sequence analysis Prior Ps and constrained Functional LRs alone, we subjected 38 variants previously classified as likely pathogenic (Class 4, n=1; Class 5, n=15) and likely neutral (Class 2, n=2; Class 1, n=20) to the two-component evaluation (Supp. Table S1). For 28 of these variants used as internal standards during assay calibration, the constrained Functional LR was derived from the LeLR calculated in the original VarCall study (Iversen et al. 2011). If two-component QIE were applied, two known pathogenic substitutions (p.R1495M and p.A1623G) would have been remanded for further analysis because their splicing Prior P was >0.05 and their splicing Prior P was greater than the Bayesian combination of the missense Prior P and the constrained Functional LR. The remaining 36 substitutions would all have been classified correctly. Including the two excluded substitutions, sensitivity was 87.5% (95% CI: 62%- 98%), specificity was 100.0% (85% -100%), and the classification rate was 94.7% (82% -99%). The error rate-classification of known neutral variants as pathogenic or vice versa – was 0.0%.

### Application to variants without patient observational data

From Woods *et al*. (2016), an additional 195 UV missense substitutions had been subjected to a functional assay and were available for two-component QIE. Of these, 36 failed the concordance test and were excluded for further analyses (Supp. Table S2). An additional 14 had splicing Prior Ps >0.05 and their splicing Prior P was greater than the Bayesian combination of the missense analysis Prior P and the constrained Functional LR; these were also remanded for further analyses (Supp. Table S2). Of the remaining 145 missense substitutions, 39 had two-component QIE Posterior Ps >0.95 and were classified as Class 4, Likely Pathogenic; 4 had Posterior P ≥0.05 and <0.95 and were classified as Class 3, Uncertain, and 102 had Posterior Ps <0.05 and were classified as Class 2, Likely Not Pathogenic.

Since there was no evidence for error in the two-component QIE analysis of known neutral or pathogenic missense substitutions, we explored the sensitivity of two-component QIE to misspecification of either the missense analysis Prior Ps or the constrained Functional LRs. For the 40 likely pathogenic substitutions (including *BRCA1* RING domain substitution p.P34S), the estimated error rate was 0.0038. Testing the consequence of the possibility that the missense analysis Prior Ps are over-estimated, lowering the Align-GVGD prior probabilities from their published point-estimates (0.81, 0.66, and 0.29 for C65, C55 to C35, and C25 & C15, respectively) (Tavtigian et al. 2008) to the published 95% lower bound of their confidence intervals (0.61, 0.34, and 0.09, in the same order) increased the expected error rate to 0.0113. As a second alternative, the constrained Functional LRs could be inflated. Testing the consequence of this possibility, we estimated the error rate as a function of constrained Functional LR deflation (Fig. 4). The constrained Functional LR would have to have been inflated by more than a factor of 5 for the error rate to reach 0.025, and buy more than a factor of 15 for the error rate to reach 0.05 (Fig. 4A). As a third alternative, the missense analysis Prior P and constrained Functional LR could be partially non-independent, so combining their full magnitude via Bayes’ rule might over-estimate the Posterior P. Since the missense analysis Prior Ps were calibrated several years before the functional assays, this was tested by reducing the magnitude of the Functional LR used to calculate the Posterior P. The linear deflation described above was one test. As the Functional LRs were reported in Woods *et al*. (2016) as L_e_LR, we also tested linear deflation of the constrained L_e_LR (Fig. 4B). The constrained Functional LRs would have to have been inflated to the 1.9^th^ power to produce an error rate of ~0.025 and to the 2.7^th^ power for the error rate to exceed 0.05.

**Figure 4.**
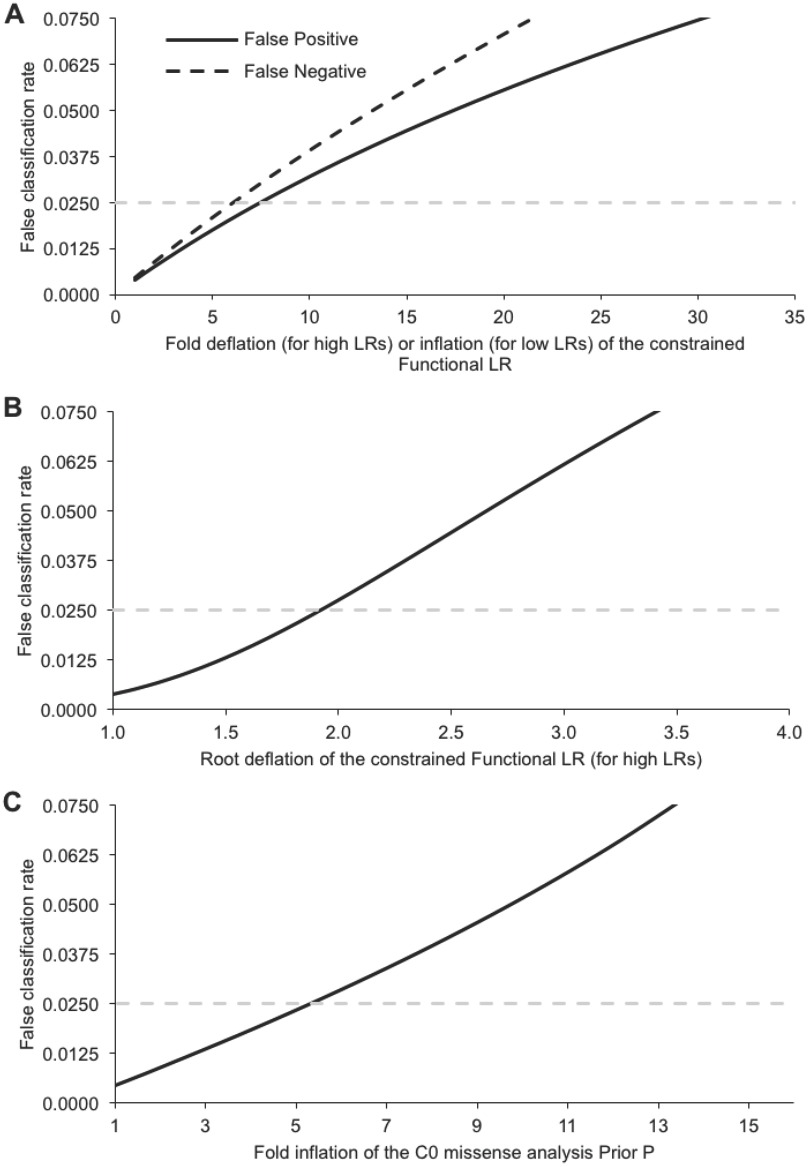
Sensitivity of two-component classification to mis-calibration of the sequence analysis-based prior probabilities or constrained Functional LR. A: Sensitivity to overestimated constrained Functional LRs for key domain missense substitutions if the missense analysis Prior Ps for the substitutions are >0.18 (solid curve), and sensitivity to underestimation of the constrained Functional LRs for key domain missense substitutions if the missense analysis Prior Ps for the substitutions are <0.05 (dashed curve). B: Sensitivity to non-independence between the missense analysis Prior Ps and constrained Functional LRs. C: Sensitivity to underestimated missense analysis Prior Ps for key domain Align-GVGD C0 missense substitutions. In each panel, the horizontal dashed grey line marks a false classification rate of 0.025.

The 102 likely neutral substitutions break into two sub-groups: 72 falling outside of the BRCT1-linker-BRCT2 key domain, and 30 falling within the key domain (see Fig. 3). The 70 upstream substitutions fall in an interval where there is no evidence for pathogenic missense substitutions except for those that damage mRNA splicing, so we did not perform any analysis. The 30 in-domain substitutions have an estimated error rate of 0.0044. These are all Align-GVGD C0 substitutions with missense analysis Prior Ps of 0.03. Testing the possibility that this missense analysis Prior P is under-estimated, increasing it to the published 95% upper bound of the confidence interval (0.06) increased the expected error rate to 0.009. If the actual Prior P was 0.15, then the estimated error rate would be ~0.025 (Fig. 4C), and if the prior was 0.30 the error rate would be ~0.05. Alternatively, the Functional LRs could be deflated. Testing the consequence of this possibility, we estimated the error rate as a function of constrained Functional LR reflation (Fig. 4A). The constrained Functional LR would have to have been deflated by more than a factor of 5 for the error rate to reach 0.025, and buy more than a factor of 10 for the error rate to reach 0.05. Since the missense analysis Prior P point estimate for these substitutions is below 0.05, and almost all of the relevant constrained Functional LRs are <1.0, linear inflation of the L_e_LR, simulating non-independence between the missense anlaysis Prior P and the constrained Functional LR does not model a false classification rate.

### Towards en masse classification

None of the 84 *BRCA1* missense substitutions between the coiled-coil and BRCT domains (aa 1,425-1,649) demonstrated loss of function in the TA assay (see Fig. 3). Using personal and summary family history data, it has previously been shown that non-spliceogenic missense substitutions falling outside of the key functional domains are associated with <2% probability of pathogenicity (Easton et al. 2007; Tavtigian et al. 2008; Vallée et al. 2016). Those results, which were based on patient observational data, are further supported by the functional assay data. Thus, one could begin to argue that all 1,410 non-spliceogenic (splicing Prior P <0.05) missense substitutions reachable by a single nucleotide substitution within this 225 amino acid segment could be *a priori* classified as Class 2, Likely Not Pathogenic.

Recently, an alternative probabilistic functional assay classification framework has been published called the ProClass toolbox (Thouvenot et al. 2016). We attempted to compare the two models using the raw TA assay data reported in the supplementary material of Woods *et al.* (2016). However, we found that ProClass in its current form could not utilize the full experimental dataset, because it could not cope with large variation between experiments for variants that had been assayed in multiple batches. Thus, we opted to use the L_e_LR generated by VarCall.

In closing, classification of sequence variants for purposes of clinical cancer genetics and patient management does not require recently deceased political commentator John McGlaughlin’s notion of “metaphysical certitude”. Accepting a “Likely Pathogenic” threshold of 0.95 implies that ~2.5% of variants so classified will turn out to have been wrongly classified. The ACMG Likely Pathogenic threshold of 0.90 (Richards et al. 2015) implies that ~5% of variants so classified will turn out to have been wrongly classified. Although the assigned ceiling, floor, and offset around L_e_LR = 0 to constrain the Functional LR are somewhat *ad hoc.* The appropriate comparators are the level of *ad hocness* in the way that the ACMG guidelines incorporates functional assays (Richards et al. 2015), and the complexity of the functional assay rules in the InSiGHT mismatch repair gene variant classification criteria (Thompson et al. 2013).

If we look back on our results 10 years from now and find that >2.5% of Likely Pathogenic variants were actually neutral, it will mean that these methods were mis-calibrated and insufficiently stringent. But if we look back 10 years from now and find that <1.0% of Likely Pathogenic variants were actually neutral, it will mean that the classification was too stringent-in fact equivalent to the Definitely Pathogenic criterion. It will also mean that fewer UVs were classified than could have been, and that fewer patients and their at-risk relatives benefited from genotype-based genetic counseling than could have been the case. The classification error rate observed when two-component QIE was applied to known neutral and known pathogenic *BRCA1* missense substitutions was 0.00. In addition, the two-component classification model appears to be reasonably robust to systematic errors in the form of (1) miscalibration of the sequence analysis-based prior probabilities of pathogenicity, (2) overly optimistic calibration of the functional assays to generate their Functional LRs, or (3) partial non-independence between the sequence analysis-based prior probabilities and the constrained Functional LRs. Therefore, it appears that we are poised to move bravely into a world of higher efficiency *BRCA1* sequence variant classification.

**Table 1.** Criteria for constraining the log_e_ likelihood ratio (L_e_LR)

| Criterion | Constrained $L_eLR$ |
| --- | --- |
| $L_eLR < -6.6$ | -4.6 |
| $-6.6 \leq L_eLR < -2.0$ | $L_eLR + 2$ |
| $-2.0 \leq L_eLR \leq 2.0$ | 0.0 |
| $2.0 < L_eLR \leq 8.9$ | $L_eLR - 2$ |
| $L_eLR > 8.9$ | 6.9 |

## Acknowledgements

BAT is a National Health and Medical Research Council CJ Martin Early Career Fellow. SVT is supported in part by the US NCI Cancer Center Support Grant P30 CA042014. Support was also provided in the form of a gift from Astra Zeneca via Newcastle University, UK, as part of the BRCA Challenge, a flagship project of the Global Alliance for Genomics and Health (GA4GH).

---

**Supplementary Table S1.** *BRCA1* C-terminus variants previously classified on Ex-UV database (n=38)

| Variant | HGVSC | HGVSP | IARC Class on Ex-UV database (Dec 2016) | Obs LR<sup>a</sup> | Obs L<sub>c</sub>LR | Status in calibration<sup>b</sup> | Functional L<sub>c</sub>LR | Constrained-L<sub>c</sub>LR |
| --- | --- | --- | --- | --- | --- | --- | --- | --- |
| H1402Y | c.4204C>T | p.H1402Y | 1 - Not pathogenic | 0.006 | -5.13 | Neut | -5.73 | -4.61 |
| E1419Q | c.4255G>C | p.E1419Q | 1 - Not pathogenic | 0.005 | -5.32 | VUS | -5.34 | -4.61 |
| N1468H | c.4402A>C | p.N1468H | 1 - Not pathogenic | 0.003 | -5.83 | VUS | -5.19 | -4.61 |
| R1495M | c.4484G>T | p.R1495M | 5 - Definitely pathogenic | 1.23x10<sup>6</sup> | 14.02 | Neut | -6.24 | NA<sup>c</sup> |
| S1512I | c.4535G>T | p.S1512I | 1 - Not pathogenic | 1.00x10<sup>-10</sup> | -23.03 | Neut | -6.62 | -4.61 |
| V1534M | c.4600G>A | p.V1534M | 1 - Not pathogenic | 0.005 | -5.34 | Neut | -6.81 | -4.61 |
| D1546N | c.4636G>A | p.D1546N | 1 - Not pathogenic | 0.006 | -5.18 | Neut | -6.51 | -4.61 |
| D1546Y | c.4636G>T | p.D1546Y | 1 - Not pathogenic | 9.30x10<sup>-4</sup> | -6.98 | VUS | -5.14 | -4.61 |
| L1564P | c.4691T>C | p.L1564P | 1 - Not pathogenic | 0.003 | -5.71 | Neut | -5.14 | -4.61 |
| S1613G | c.4837A>G | p.S1613G | 1 - Not pathogenic | 1.00x10<sup>-10</sup> | -23.03 | Neut | -7.03 | -4.61 |
| P1614S | c.4840C>T | p.P1614S | 1 - Not pathogenic | 2.24x10<sup>-9</sup> | -19.92 | Neut | -6.82 | -4.61 |
| A1623G | c.4868C>G | p.A1623G | 5 - Definitely pathogenic | 389.1 | 5.96 | VUS | -5.16 | NA<sup>c</sup> |
| M1628T | c.4883T>C | p.M1628T | 1 - Not pathogenic | 1.60x10<sup>-4</sup> | -8.74 | Neut | -5.77 | -4.61 |
| P1637L | c.4910C>T | p.P1637L | 1 - Not pathogenic | 0.002 | -6.19 | VUS | -5.06 | -4.61 |
| M1652T | c.4955T>C | p.M1652T | 1 - Not pathogenic | 2.70x10<sup>-4</sup> | -8.22 | Neut | NA | NA |
| M1652I | c.4956G>A | p.M1652I | 1 - Not pathogenic | 4.80x10<sup>-4</sup> | -7.64 | Neut | -6.63 | -4.61 |
| F1662S | c.4985T>C | p.F1662S | 1 - Not pathogenic | 0.005 | -5.28 | Neut | -6.40 | -4.61 |
| L1664P | c.4991T>C | p.L1664P | 1 - Not pathogenic | 3.87x10<sup>-4</sup> | -7.86 | Neut | -6.38 | -4.61 |
| E1682K | c.5044G>A | p.E1682K | 1 - Not pathogenic | 0.003 | -5.76 | Neut | NA | NA |
| T1685A | c.5053A>G | p.T1685A | 5 - Definitely pathogenic | 107.0 | 4.67 | Path | NA | NA |
| T1685I | c.5054C>T | p.T1685I | 5 - Definitely pathogenic | 138.5 | 4.93 | Path | 18.59 | 6.91 |
| M1689R | c.5066T>G | p.M1689R | 4 - Likely pathogenic | 45.73 | 3.82 | Path | 16.00 | 6.91 |
| R1699W | c.5095C>T | p.R1699W | 5 - Definitely pathogenic | 3.98x10<sup>4</sup> | 10.59 | Path | 1.99 | 0.00 |
| R1699Q | c.5096G>A | p.R1699Q | 3 - Uncertain | 3.976 | 1.38 | Path | 6.65 | 4.65 |
| G1706E | c.5117G>A | p.G1706E | 5 - Definitely pathogenic | 589.3 | 6.38 | Path | 16.61 | 6.91 |
| G1706A | c.5117G>C | p.G1706A | 1 - Not pathogenic | 2.55x10<sup>-5</sup> | -10.58 | Neut | NA | NA |
| A1708E | c.5123C>A | p.A1708E | 5 - Definitely pathogenic | 2.09x10<sup>12</sup> | 28.37 | Path | 17.67 | 6.91 |
| A1708V | c.5123C>T | p.A1708V | 3 - Uncertain | 0.044 | -3.13 | VUS | 2.66 | 0.66 |
| S1715R | c.5143A>C | p.S1715R | 5 - Definitely pathogenic | 44.66 | 3.80 | Path | 8.16 | 6.91 |
| T1720A | c.5158A>G | p.T1720A | 1 - Not pathogenic | 1.82x10<sup>-11</sup> | -24.73 | Neut | -6.59 | -4.61 |
| V1736A | c.5207T>C | p.V1736A | 5 - Definitely pathogenic | 234.0 | 5.46 | VUS | 21.87 | 6.91 |
| G1738R | c.5212G>A | p.G1738R | 5 - Definitely pathogenic | 115.0 | 4.74 | Path | 17.39 | 6.91 |
| R1751Q | c.5252G>A | p.R1751Q | 1 - Not pathogenic | 0.007 | -4.99 | Neut | -6.11 | -4.61 |
| L1764P | c.5291T>C | p.L1764P | 5 - Definitely pathogenic | 346.6 | 5.85 | Path | 16.85 | 6.91 |
| I1766S | c.5297T>G | p.I1766S | 5 - Definitely pathogenic | 138.4 | 4.93 | Path | 16.01 | 6.91 |
| M1775K | c.5324T>A | p.M1775K | 5 - Definitely pathogenic | 235.0 | 5.46 | Path | 13.37 | 6.91 |
| M1775R | c.5324T>G | p.M1775R | 5 - Definitely pathogenic | 229.0 | 5.43 | Path | 17.24 | 6.91 |
| P1776H | c.5327C>A | p.P1776H | 2 - Likely not pathogenic | 1.039 | 0.04 | VUS | -5.30 | -4.61 |
| D1778G | c.5333A>G | p.D1778G | 2 - Likely not pathogenic | 0.330 | -1.11 | VUS | -3.29 | -1.29 |
| C1787S/<br>G1788D<sup>d</sup> | c.[5359T>A;<br>5363G>A] | p.[C1787S;<br>G1788D] | 5 - Definitely pathogenic | 2032.8 | 7.62 | VUS | 30.76 | 6.91 |
| G1788V | c.5363G>T | p.G1788V | 5 - Definitely pathogenic | 7077.7 | 8.86 | Path | 17.80 | 6.91 |
| V1804D | c.5411T>A | p.V1804D | 1 - Not pathogenic | 0.001 | -6.73 | Neut | -5.17 | -4.61 |
| V1838E | c.5513T>A | p.V1838E | 5 - Definitely pathogenic | 3974.0 | 8.29 | Path | NA | NA |
| I1858L | c.5572A>C | p.I1858L | 1 - Not pathogenic | 0.002 | -6.08 | VUS | -5.12 | -4.61 |
| P1859R | c.5576C>G | p.P1859R | 1 - Not pathogenic | 3.31x10<sup>-5</sup> | -10.31 | Neut | -6.64 | -4.61 |

L<sub>e</sub>LR – loge likelihood ratio
<sup>a</sup> Odds in favor of pathogenicity based on patient observational (Obs) data, equivalent to Product LR on Ex-UV database.
<sup>b</sup> Neut were assigned as neutral controls, and Path were assigned as pathogenic controls in VarCall calibration (Woods et al 2016 npj Genomic Med 1:16001). For the Path and Neut variants, the L<sub>e</sub>LR was taken from Iversen et al 2011 Cancer Epidemiol Biomarkers Prev 20:1078–1088.
<sup>c</sup> Spliceogenic pathogenic variant.
<sup>d</sup> Was originally reported in the Ex-UV database as C1787S only

**Supplementary Table S2.** Two-component quantitative integrated evaluation of *BRCA1* C-terminus variants without observational data (n=195)

| Variant | HGVSC | HGVSP | L<sub>c</sub>LR | Constrained-L<sub>c</sub>LR | Functional LR | Splicing Prior P | Missense analysis Prior P | Applicable Prior P | Posterior P | New IARC Class |
| --- | --- | --- | --- | --- | --- | --- | --- | --- | --- | --- |
| E1352K | c.4054G>A | p.E1352K | -3.13 | -1.13 | 0.32 | 0.02 | 0.02 | 0.02 | 0.007 | 2 - Likely Not Pathogenic |
| C1372Y | c.4115G>A | p.C1372Y | -4.56 | -2.56 | 0.08 | 0.3 | 0.02 | 0.3 | 0.002 | NA - node 4 |
| Q1395R | c.4184A>G | p.Q1395R | -3.84 | -1.84 | 0.16 | 0.34 | 0.02 | 0.34 | 0.003 | NA - node 4 |
| M1400V | c.4198A>G | p.M1400V | -4.66 | -2.66 | 0.07 | 0.02 | 0.02 | 0.02 | 0.001 | 2 - Likely Not Pathogenic |
| M1400T | c.4199T>C | p.M1400T | -4.25 | -2.25 | 0.11 | 0.02 | 0.02 | 0.02 | 0.002 | 2 - Likely Not Pathogenic |
| M1400I | c.4200G>C | p.M1400I | -4.46 | -2.46 | 0.09 | 0.02 | 0.02 | 0.02 | 0.002 | 2 - Likely Not Pathogenic |
| H1402R | c.4205A>G | p.H1402R | -3.57 | -1.57 | 0.21 | 0.3 | 0.02 | 0.3 | 0.004 | NA - node 4 |
| L1404P | c.4211T>C | p.L1404P | -1.84 | 0.00 | 1.00 | 0.02 | 0.02 | 0.02 | 0.020 | NA - node 3 |
| I1405V | c.4213A>G | p.I1405V | -4.63 | -2.63 | 0.07 | 0.3 | 0.02 | 0.3 | 0.001 | NA - node 4 |
| L1407P | c.4220T>C | p.L1407P | 2.48 | 0.48 | 1.62 | 0.02 | 0.02 | 0.02 | 0.032 | NA - node 3 |
| M1411T | c.4232T>C | p.M1411T | 1.86 | 0.00 | 1.00 | 0.02 | 0.02 | 0.02 | 0.020 | NA - node 3 |
| A1416T | c.4246G>A | p.A1416T | -3.60 | -1.60 | 0.20 | 0.02 | 0.02 | 0.02 | 0.004 | 2 - Likely Not Pathogenic |
| H1421Y | c.4261C>T | p.H1421Y | -3.77 | -1.77 | 0.17 | 0.02 | 0.02 | 0.02 | 0.003 | 2 - Likely Not Pathogenic |
| P1430S | c.4288C>T | p.P1430S | -4.99 | -2.99 | 0.05 | 0.02 | 0.02 | 0.02 | 0.001 | 2 - Likely Not Pathogenic |
| I1432L | c.4294A>C | p.I1432L | -5.28 | -3.28 | 0.04 | 0.02 | 0.02 | 0.02 | 0.001 | 2 - Likely Not Pathogenic |
| R1443Q | c.4328G>A | p.R1443Q | -5.14 | -3.14 | 0.04 | 0.02 | 0.02 | 0.02 | 0.001 | 2 - Likely Not Pathogenic |
| S1448G | c.4342A>G | p.S1448G | -5.02 | -3.02 | 0.05 | 0.02 | 0.02 | 0.02 | 0.001 | 2 - Likely Not Pathogenic |
| S1448T | c.4343G>C | p.S1448T | -5.15 | -3.15 | 0.04 | 0.02 | 0.02 | 0.02 | 0.001 | 2 - Likely Not Pathogenic |
| S1457A | c.4369T>G | p.S1457A | -4.96 | -2.96 | 0.05 | 0.02 | 0.02 | 0.02 | 0.001 | 2 - Likely Not Pathogenic |
| E1462K | c.4384G>A | p.E1462K | -4.64 | -2.64 | 0.07 | 0.02 | 0.02 | 0.02 | 0.001 | 2 - Likely Not Pathogenic |
| P1469S | c.4405C>T | p.P1469S | -5.12 | -3.12 | 0.04 | 0.02 | 0.02 | 0.02 | 0.001 | 2 - Likely Not Pathogenic |
| E1470D | c.4410A>C | p.E1470D | -5.23 | -3.23 | 0.04 | 0.02 | 0.02 | 0.02 | 0.001 | 2 - Likely Not Pathogenic |
| S1484T | c.4450T>A | p.S1484T | -5.22 | -3.22 | 0.04 | 0.02 | 0.02 | 0.02 | 0.001 | 2 - Likely Not Pathogenic |
| T1485I | c.4454C>T | p.T1485I | -5.27 | -3.27 | 0.04 | 0.02 | 0.02 | 0.02 | 0.001 | 2 - Likely Not Pathogenic |
| S1486C | c.4456A>T | p.S1486C | -5.15 | -3.15 | 0.04 | 0.02 | 0.02 | 0.02 | 0.001 | 2 - Likely Not Pathogenic |
| K1487R | c.4460A>G | p.K1487R | -4.14 | -2.14 | 0.12 | 0.02 | 0.02 | 0.02 | 0.002 | 2 - Likely Not Pathogenic |
| E1494K | c.4480G>A | p.E1494K | -4.61 | -2.61 | 0.07 | 0.02 | 0.02 | 0.02 | 0.001 | 2 - Likely Not Pathogenic |
| R1495K | c.4484G>A | p.R1495K | -4.94 | -2.94 | 0.05 | 0.34 | 0.02 | 0.34 | 0.001 | NA - node 4 |
| S1497A | c.4489T>G | p.S1497A | -5.05 | -3.05 | 0.05 | 0.3 | 0.02 | 0.3 | 0.001 | NA - node 4 |
| P1502S | c.4504C>T | p.P1502S | -5.12 | -3.12 | 0.04 | 0.02 | 0.02 | 0.02 | 0.001 | 2 - Likely Not Pathogenic |
| R1507T | c.4520G>C | p.R1507T | -5.32 | -3.32 | 0.04 | 0.02 | 0.02 | 0.02 | 0.001 | 2 - Likely Not Pathogenic |
| S1512C | c.4534A>T | p.S1512C | -5.31 | -3.31 | 0.04 | 0.02 | 0.02 | 0.02 | 0.001 | 2 - Likely Not Pathogenic |
| L1517F | c.4549C>T | p.L1517F | -5.31 | -3.31 | 0.04 | 0.02 | 0.02 | 0.02 | 0.001 | 2 - Likely Not Pathogenic |
| Y1522C | c.4565A>G | p.Y1522C | -4.65 | -2.65 | 0.07 | 0.02 | 0.02 | 0.02 | 0.001 | 2 - Likely Not Pathogenic |
| S1524A | c.4570T>G | p.S1524A | -5.21 | -3.21 | 0.04 | 0.02 | 0.02 | 0.02 | 0.001 | 2 - Likely Not Pathogenic |
| E1527K | c.4579G>A | p.E1527K | -4.49 | -2.49 | 0.08 | 0.02 | 0.02 | 0.02 | 0.002 | 2 - Likely Not Pathogenic |
| I1529V | c.4585A>G | p.I1529V | -4.92 | -2.92 | 0.05 | 0.02 | 0.02 | 0.02 | 0.001 | 2 - Likely Not Pathogenic |
| L1539P | c.4616T>C | p.L1539P | -4.15 | -2.15 | 0.12 | 0.02 | 0.02 | 0.02 | 0.002 | 2 - Likely Not Pathogenic |
| S1542A | c.4624T>G | p.S1542A | -5.27 | -3.27 | 0.04 | 0.02 | 0.02 | 0.02 | 0.001 | 2 - Likely Not Pathogenic |
| S1542C | c.4625C>G | p.S1542C | -5.10 | -3.10 | 0.04 | 0.02 | 0.02 | 0.02 | 0.001 | 2 - Likely Not Pathogenic |
| P1544L | c.4631C>T | p.P1544L | -5.32 | -3.32 | 0.04 | 0.02 | 0.02 | 0.02 | 0.001 | 2 - Likely Not Pathogenic |
| T1548M | c.4643C>T | p.T1548M | -5.19 | -3.19 | 0.04 | 0.02 | 0.02 | 0.02 | 0.001 | 2 - Likely Not Pathogenic |
| T1550I | c.4649C>T | p.T1550I | -5.27 | -3.27 | 0.04 | 0.02 | 0.02 | 0.02 | 0.001 | 2 - Likely Not Pathogenic |
| L1553M | c.4657T>A | p.L1553M | -4.70 | -2.70 | 0.07 | 0.02 | 0.02 | 0.02 | 0.001 | 2 - Likely Not Pathogenic |
| D1557H | c.4669G>C | p.D1557H | -4.85 | -2.85 | 0.06 | 0.02 | 0.02 | 0.02 | 0.001 | 2 - Likely Not Pathogenic |
| E1559K | c.4675G>A | p.E1559K | -3.71 | -1.71 | 0.18 | 0.97 | 0.02 | 0.97 | 0.004 | NA - node 4 |
| E1559Q | c.4675G>C | p.E1559Q | -4.69 | -2.69 | 0.07 | 0.97 | 0.02 | 0.97 | 0.001 | NA - node 4 |
| T1561I | c.4682C>T | p.T1561I | -5.26 | -3.26 | 0.04 | 0.02 | 0.02 | 0.02 | 0.001 | 2 - Likely Not Pathogenic |
| P1562L | c.4685C>T | p.P1562L | -5.14 | -3.14 | 0.04 | 0.02 | 0.02 | 0.02 | 0.001 | 2 - Likely Not Pathogenic |
| I1568V | c.4702A>G | p.I1568V | -4.15 | -2.15 | 0.12 | 0.02 | 0.02 | 0.02 | 0.002 | 2 - Likely Not Pathogenic |
| L1570F | c.4708C>T | p.L1570F | -5.21 | -3.21 | 0.04 | 0.02 | 0.02 | 0.02 | 0.001 | 2 - Likely Not Pathogenic |
| F1571S | c.4712T>C | p.F1571S | -2.35 | -0.35 | 0.70 | 0.02 | 0.02 | 0.02 | 0.014 | 2 - Likely Not Pathogenic |
| P1575H | c.4724C>A | p.P1575H | -4.63 | -2.63 | 0.07 | 0.02 | 0.02 | 0.02 | 0.001 | 2 - Likely Not Pathogenic |
| S1577P | c.4729T>C | p.S1577P | -5.01 | -3.01 | 0.05 | 0.02 | 0.02 | 0.02 | 0.001 | 2 - Likely Not Pathogenic |
| D1578G | c.4733A>G | p.D1578G | -4.81 | -2.81 | 0.06 | 0.02 | 0.02 | 0.02 | 0.001 | 2 - Likely Not Pathogenic |
| P1579A | c.4735C>G | p.P1579A | -4.74 | -2.74 | 0.06 | 0.02 | 0.02 | 0.02 | 0.001 | 2 - Likely Not Pathogenic |
| S1580Y | c.4739C>A | p.S1580Y | -4.47 | -2.47 | 0.08 | 0.02 | 0.02 | 0.02 | 0.002 | 2 - Likely Not Pathogenic |
| S1580F | c.4739C>T | p.S1580F | -5.25 | -3.25 | 0.04 | 0.02 | 0.02 | 0.02 | 0.001 | 2 - Likely Not Pathogenic |
| A1584S | c.4750G>T | p.A1584S | -4.90 | -2.90 | 0.05 | 0.02 | 0.02 | 0.02 | 0.001 | 2 - Likely Not Pathogenic |
| R1589C | c.4765C>T | p.R1589C | -5.23 | -3.23 | 0.04 | 0.02 | 0.02 | 0.02 | 0.001 | 2 - Likely Not Pathogenic |
| R1589H | c.4766G>A | p.R1589H | -5.08 | -3.08 | 0.05 | 0.02 | 0.02 | 0.02 | 0.001 | 2 - Likely Not Pathogenic |
| S1596L | c.4787C>T | p.S1596L | -5.17 | -3.17 | 0.04 | 0.02 | 0.02 | 0.02 | 0.001 | 2 - Likely Not Pathogenic |
| T1597A | c.4789A>G | p.T1597A | -4.84 | -2.84 | 0.06 | 0.02 | 0.02 | 0.02 | 0.001 | 2 - Likely Not Pathogenic |
| L1600F | c.4800G>C | p.L1600F | -4.73 | -2.73 | 0.07 | 0.02 | 0.02 | 0.02 | 0.001 | 2 - Likely Not Pathogenic |
| K1601E | c.4801A>G | p.K1601E | -5.17 | -3.17 | 0.04 | 0.02 | 0.02 | 0.02 | 0.001 | 2 - Likely Not Pathogenic |
| Q1604R | c.4811A>G | p.Q1604R | -4.67 | -2.67 | 0.07 | 0.02 | 0.02 | 0.02 | 0.001 | 2 - Likely Not Pathogenic |
| L1605V | c.4813T>G | p.L1605V | -5.02 | -3.02 | 0.05 | 0.02 | 0.02 | 0.02 | 0.001 | 2 - Likely Not Pathogenic |
| K1606E | c.4816A>G | p.K1606E | -5.38 | -3.38 | 0.03 | 0.02 | 0.02 | 0.02 | 0.001 | 2 - Likely Not Pathogenic |
| A1608V | c.4823C>T | p.A1608V | -5.02 | -3.02 | 0.05 | 0.02 | 0.02 | 0.02 | 0.001 | 2 - Likely Not Pathogenic |
| S1613C | c.4837A>T | p.S1613C | -5.07 | -3.07 | 0.05 | 0.02 | 0.02 | 0.02 | 0.001 | 2 - Likely Not Pathogenic |
| A1615T | c.4843G>A | p.A1615T | -5.18 | -3.18 | 0.04 | 0.02 | 0.02 | 0.02 | 0.001 | 2 - Likely Not Pathogenic |
| A1615D | c.4844C>A | p.A1615D | -4.81 | -2.81 | 0.06 | 0.02 | 0.02 | 0.02 | 0.001 | 2 - Likely Not Pathogenic |
| M1628V | c.4882A>G | p.M1628V | -4.20 | -2.20 | 0.11 | 0.02 | 0.02 | 0.02 | 0.002 | 2 - Likely Not Pathogenic |
| M1628I | c.4884G>C | p.M1628I | -5.20 | -3.20 | 0.04 | 0.02 | 0.02 | 0.02 | 0.001 | 2 - Likely Not Pathogenic |
| S1631N | c.4892G>A | p.S1631N | -5.29 | -3.29 | 0.04 | 0.3 | 0.02 | 0.3 | 0.001 | NA - node 4 |
| A1641T | c.4921G>A | p.A1641T | -5.01 | -3.01 | 0.05 | 0.02 | 0.02 | 0.02 | 0.001 | 2 - Likely Not Pathogenic |
| E1644G | c.4931A>G | p.E1644G | -5.00 | -3.00 | 0.05 | 0.02 | 0.02 | 0.02 | 0.001 | 2 - Likely Not Pathogenic |
| R1645G | c.4933A>G | p.R1645G | -5.26 | -3.26 | 0.04 | 0.02 | 0.02 | 0.02 | 0.001 | 2 - Likely Not Pathogenic |
| R1645T | c.4934G>C | p.R1645T | -5.23 | -3.23 | 0.04 | 0.02 | 0.02 | 0.02 | 0.001 | 2 - Likely Not Pathogenic |
| R1645M | c.4934G>T | p.R1645M | -5.24 | -3.24 | 0.04 | 0.02 | 0.02 | 0.02 | 0.001 | 2 - Likely Not Pathogenic |
| R1645S | c.4935G>C | p.R1645S | -5.14 | -3.14 | 0.04 | 0.02 | 0.02 | 0.02 | 0.001 | 2 - Likely Not Pathogenic |
| N1647K | c.4941C>A | p.N1647K | -4.56 | -2.56 | 0.08 | 0.02 | 0.02 | 0.02 | 0.002 | 2 - Likely Not Pathogenic |
| S1651F | c.4952C>T | p.S1651F | -0.57 | 0.00 | 1.00 | 0.02 | 0.03 | 0.03 | 0.030 | NA - node 3 |
| M1652K | c.4955T>A | p.M1652K | 32.95 | 6.91 | 1000 | 0.02 | 0.66 | 0.66 | 0.999 | 4 - Likely Pathogenic |
| V1653M | c.4957G>A | p.V1653M | 8.64 | 6.64 | 762.6 | 0.02 | 0.03 | 0.03 | 0.959 | NA - node 3 |
| S1655F | c.4964C>T | p.S1655F | 14.00 | 6.91 | 1000 | 0.02 | 0.29 | 0.29 | 0.998 | 4 - Likely Pathogenic |
| G1656D | c.4967G>A | p.G1656D | 0.55 | 0.00 | 1.00 | 0.02 | 0.81 | 0.81 | 0.810 | NA - node 3 |
| E1660V | c.4979A>T | p.E1660V | -5.55 | -3.55 | 0.03 | 0.02 | 0.03 | 0.03 | 0.001 | 2 - Likely Not Pathogenic |
| M1663L | c.4987A>C | p.M1663L | -2.53 | -0.53 | 0.59 | 0.34 | 0.03 | 0.34 | 0.018 | NA - node 4 |
| M1663K | c.4988T>A | p.M1663K | -3.64 | -1.64 | 0.19 | 0.34 | 0.03 | 0.34 | 0.006 | NA - node 4 |
| V1665M | c.4993G>A | p.V1665M | -1.29 | 0.00 | 1.00 | 0.02 | 0.29 | 0.29 | 0.290 | NA - node 3 |
| A1669S | c.5005G>T | p.A1669S | -0.77 | 0.00 | 1.00 | 0.02 | 0.03 | 0.03 | 0.030 | NA - node 3 |
| L1679Q | c.5036T>A | p.L1679Q | -5.60 | -3.60 | 0.03 | 0.02 | 0.03 | 0.03 | 0.001 | 2 - Likely Not Pathogenic |
| E1682V | c.5045A>T | p.E1682V | -4.34 | -2.34 | 0.10 | 0.02 | 0.03 | 0.03 | 0.003 | 2 - Likely Not Pathogenic |
| M1689T | c.5066T>C | p.M1689T | 0.09 | 0.00 | 1.00 | 0.02 | 0.29 | 0.29 | 0.290 | NA - node 3 |
| T1691K | c.5072C>A | p.T1691K | 34.17 | 6.91 | 1000 | 0.04 | 0.81 | 0.81 | 1.000 | 4 - Likely Pathogenic |
| T1691I | c.5072C>T | p.T1691I | 12.84 | 6.91 | 1000 | 0.34 | 0.81 | 0.81 | 1.000 | 4 - Likely Pathogenic |
| D1692N | c.5074G>A | p.D1692N | -1.54 | 0.00 | 1.00 | 0.97 | 0.29 | 0.97 | 0.290 | NA - node 3 |
| D1692H | c.5074G>C | p.D1692H | 13.86 | 6.91 | 1000 | 0.97 | 0.81 | 0.97 | 1.000 | 4 - Likely Pathogenic |
| D1692Y | c.5074G>T | p.D1692Y | 7.45 | 5.45 | 233.0 | 0.97 | 0.81 | 0.97 | 0.999 | 4 - Likely Pathogenic |
| F1695L | c.5083T>C | p.F1695L | -2.62 | -0.62 | 0.54 | 0.02 | 0.03 | 0.03 | 0.016 | 2 - Likely Not Pathogenic |
| V1696L | c.5086G>C | p.V1696L | -1.85 | 0.00 | 1.00 | 0.02 | 0.29 | 0.29 | 0.290 | NA - node 3 |
| C1697R | c.5089T>C | p.C1697R | 19.97 | 6.91 | 1000 | 0.02 | 0.81 | 0.81 | 1.000 | 4 - Likely Pathogenic |
| C1697Y | c.5090G>A | p.C1697Y | 33.27 | 6.91 | 1000 | 0.02 | 0.81 | 0.81 | 1.000 | 4 - Likely Pathogenic |
| C1697F | c.5090G>T | p.C1697F | 23.82 | 6.91 | 1000 | 0.02 | 0.81 | 0.81 | 1.000 | 4 - Likely Pathogenic |
| R1699P | c.5096G>C | p.R1699P | -2.90 | -0.90 | 0.41 | 0.02 | 0.81 | 0.81 | 0.634 | NA - node 3 |
| R1699L | c.5096G>T | p.R1699L | 21.54 | 6.91 | 1000 | 0.02 | 0.81 | 0.81 | 1.000 | 4 - Likely Pathogenic |
| T1700A | c.5098A>G | p.T1700A | 2.66 | 0.66 | 1.93 | 0.02 | 0.66 | 0.66 | 0.790 | 3 - Uncertain |
| V1713A | c.5138T>C | p.V1713A | 22.28 | 6.91 | 1000 | 0.02 | 0.29 | 0.29 | 0.998 | 4 - Likely Pathogenic |
| V1714G | c.5141T>G | p.V1714G | 4.52 | 2.52 | 12.41 | 0.3 | 0.81 | 0.81 | 0.981 | 4 - Likely Pathogenic |
| S1715C | c.5143A>T | p.S1715C | 9.37 | 6.91 | 1000 | 0.02 | 0.81 | 0.81 | 1.000 | 4 - Likely Pathogenic |
| S1715N | c.5144G>A | p.S1715N | 11.57 | 6.91 | 1000 | 0.02 | 0.66 | 0.66 | 0.999 | 4 - Likely Pathogenic |
| W1718S | c.5153G>C | p.W1718S | 6.21 | 4.21 | 67.43 | 0.34 | 0.81 | 0.81 | 0.997 | 4 - Likely Pathogenic |
| W1718C | c.5154G>C | p.W1718C | 18.70 | 6.91 | 1000 | 0.34 | 0.81 | 0.81 | 1.000 | 4 - Likely Pathogenic |
| T1720I | c.5159C>T | p.T1720I | -5.09 | -3.09 | 0.05 | 0.02 | 0.03 | 0.03 | 0.001 | 2 - Likely Not Pathogenic |
| S1722F | c.5165C>T | p.S1722F | 10.12 | 6.91 | 1000 | 0.02 | 0.29 | 0.29 | 0.998 | 4 - Likely Pathogenic |
| R1726G | c.5176A>G | p.R1726G | -0.18 | 0.00 | 1.00 | 0.02 | 0.03 | 0.03 | 0.030 | NA - node 3 |
| N1730S | c.5189A>G | p.N1730S | -3.85 | -1.85 | 0.16 | 0.02 | 0.03 | 0.03 | 0.005 | 2 - Likely Not Pathogenic |
| E1731D | c.5193G>C | p.E1731D | -4.84 | -2.84 | 0.06 | 0.04 | 0.03 | 0.03 | 0.002 | 2 - Likely Not Pathogenic |
| D1733Y | c.5197G>T | p.D1733Y | -5.40 | -3.40 | 0.03 | 0.02 | 0.03 | 0.03 | 0.001 | 2 - Likely Not Pathogenic |
| D1733G | c.5198A>G | p.D1733G | -3.40 | -1.40 | 0.25 | 0.02 | 0.03 | 0.03 | 0.008 | 2 - Likely Not Pathogenic |
| F1734S | c.5201T>C | p.F1734S | -3.05 | -1.05 | 0.35 | 0.02 | 0.66 | 0.66 | 0.406 | NA - node 3 |
| V1736G | c.5207T>G | p.V1736G | 10.21 | 6.91 | 1000 | 0.02 | 0.81 | 0.81 | 1.000 | 4 - Likely Pathogenic |
| G1738E | c.5213G>A | p.G1738E | 13.65 | 6.91 | 1000 | 0.02 | 0.81 | 0.81 | 1.000 | 4 - Likely Pathogenic |
| D1739Y | c.5215G>T | p.D1739Y | 12.01 | 6.91 | 1000 | 0.02 | 0.81 | 0.81 | 1.000 | 4 - Likely Pathogenic |
| D1739G | c.5216A>G | p.D1739G | 5.81 | 3.81 | 45.12 | 0.3 | 0.81 | 0.81 | 0.995 | 4 - Likely Pathogenic |
| D1739V | c.5216A>T | p.D1739V | 5.88 | 3.88 | 48.30 | 0.02 | 0.81 | 0.81 | 0.995 | 4 - Likely Pathogenic |
| D1739E | c.5217T>G | p.D1739E | 10.95 | 6.91 | 1000 | 0.02 | 0.66 | 0.66 | 0.999 | 4 - Likely Pathogenic |
| V1741G | c.5222T>G | p.V1741G | 4.60 | 2.60 | 13.51 | 0.02 | 0.66 | 0.66 | 0.963 | 4 - Likely Pathogenic |
| H1746N | c.5236C>A | p.H1746N | -0.42 | 0.00 | 1.00 | 0.02 | 0.81 | 0.81 | 0.810 | NA - node 3 |
| H1746Y | c.5236C>T | p.H1746Y | 1.59 | 0.00 | 1.00 | 0.02 | 0.81 | 0.81 | 0.810 | NA - node 3 |
| G1748D | c.5243G>A | p.G1748D | 30.09 | 6.91 | 1000 | 0.02 | 0.81 | 0.81 | 1.000 | 4 - Likely Pathogenic |
| P1749R | c.5246C>G | p.P1749R | 13.01 | 6.91 | 1000 | 0.02 | 0.81 | 0.81 | 1.000 | 4 - Likely Pathogenic |
| R1751P | c.5252G>C | p.R1751P | 28.90 | 6.91 | 1000 | 0.02 | 0.66 | 0.66 | 0.999 | 4 - Likely Pathogenic |
| A1752T | c.5254G>A | p.A1752T | 17.90 | 6.91 | 1000 | 0.02 | 0.03 | 0.03 | 0.969 | NA - node 3 |
| A1752P | c.5254G>C | p.A1752P | 25.18 | 6.91 | 1000 | 0.02 | 0.03 | 0.03 | 0.969 | NA - node 3 |
| A1752V | c.5255C>T | p.A1752V | 4.35 | 2.35 | 10.47 | 0.3 | 0.03 | 0.3 | 0.245 | NA - node 3 |
| R1753T | c.5258G>C | p.R1753T | 21.77 | 6.91 | 1000 | 0.02 | 0.81 | 0.81 | 1.000 | 4 - Likely Pathogenic |
| E1754V | c.5261A>T | p.E1754V | -5.26 | -3.26 | 0.04 | 0.02 | 0.03 | 0.03 | 0.001 | 2 - Likely Not Pathogenic |
| F1761S | c.5282T>C | p.F1761S | 8.43 | 6.43 | 622.9 | 0.02 | 0.66 | 0.66 | 0.999 | 4 - Likely Pathogenic |
| G1763V | c.5288G>T | p.G1763V | 2.32 | 0.32 | 1.38 | 0.02 | 0.03 | 0.03 | 0.041 | 2 - Likely Not Pathogenic |
| E1765K | c.5293G>A | p.E1765K | -5.42 | -3.42 | 0.03 | 0.02 | 0.03 | 0.03 | 0.001 | 2 - Likely Not Pathogenic |
| P1771R | c.5312C>G | p.P1771R | -3.21 | -1.21 | 0.30 | 0.02 | 0.29 | 0.29 | 0.108 | NA - node 3 |
| P1771L | c.5312C>T | p.P1771L | -2.47 | -0.47 | 0.63 | 0.02 | 0.29 | 0.29 | 0.204 | 3 - Uncertain |
| T1773S | c.5317A>T | p.T1773S | -3.14 | -1.14 | 0.32 | 0.02 | 0.66 | 0.66 | 0.382 | NA - node 3 |
| T1773I | c.5318C>T | p.T1773I | -2.11 | -0.11 | 0.89 | 0.02 | 0.81 | 0.81 | 0.792 | 3 - Uncertain |
| D1778N | c.5332G>A | p.D1778N | -3.12 | -1.12 | 0.33 | 0.34 | 0.03 | 0.34 | 0.010 | NA - node 4 |
| D1778Y | c.5332G>T | p.D1778Y | -2.82 | -0.82 | 0.44 | 0.34 | 0.29 | 0.34 | 0.152 | NA - node 3 |
| L1780P | c.5339T>C | p.L1780P | 9.79 | 6.91 | 1000 | 0.02 | 0.66 | 0.66 | 0.999 | 4 - Likely Pathogenic |
| M1783L | c.5347A>C | p.M1783L | -4.40 | -2.40 | 0.09 | 0.3 | 0.03 | 0.3 | 0.003 | NA - node 4 |
| M1783T | c.5348T>C | p.M1783T | -1.40 | 0.00 | 1.00 | 0.3 | 0.66 | 0.66 | 0.660 | NA - node 3 |
| M1783I | c.5349G>C | p.M1783I | -3.48 | -1.48 | 0.23 | 0.02 | 0.03 | 0.03 | 0.007 | 2 - Likely Not Pathogenic |
| Q1785H | c.5355G>C | p.Q1785H | 0.04 | 0.00 | 1.00 | 0.02 | 0.03 | 0.03 | 0.030 | NA - node 3 |
| C1787S | c.5359T>A | p.C1787S | -5.15 | -3.15 | 0.04 | 0.02 | 0.81 | 0.81 | 0.155 | NA - node 3 |
| G1788D | c.5363G>A | p.G1788D | -2.02 | -0.02 | 0.98 | 0.02 | 0.81 | 0.81 | 0.808 | 3 - Uncertain |
| A1789T | c.5365G>A | p.A1789T | 29.40 | 6.91 | 1000 | 0.3 | 0.66 | 0.66 | 0.999 | 4 - Likely Pathogenic |
| A1789S | c.5365G>T | p.A1789S | -3.66 | -1.66 | 0.19 | 0.02 | 0.81 | 0.81 | 0.447 | NA - node 3 |
| S1790Y | c.5369C>A | p.S1790Y | -5.18 | -3.18 | 0.04 | 0.02 | 0.29 | 0.29 | 0.017 | NA - node 3 |
| V1791L | c.5371G>C | p.V1791L | -4.66 | -2.66 | 0.07 | 0.02 | 0.03 | 0.03 | 0.002 | 2 - Likely Not Pathogenic |
| E1794D | c.5382G>C | p.E1794D | -5.06 | -3.06 | 0.05 | 0.02 | 0.03 | 0.03 | 0.001 | 2 - Likely Not Pathogenic |
| G1803A | c.5408G>C | p.G1803A | 11.34 | 6.91 | 1000 | 0.34 | 0.03 | 0.34 | 0.969 | NA - node 3 |
| V1804A | c.5411T>C | p.V1804A | -5.07 | -3.07 | 0.05 | 0.02 | 0.03 | 0.03 | 0.001 | 2 - Likely Not Pathogenic |
| H1805P | c.5414A>C | p.H1805P | -4.03 | -2.03 | 0.13 | 0.02 | 0.03 | 0.03 | 0.004 | 2 - Likely Not Pathogenic |
| H1805L | c.5414A>T | p.H1805L | -5.40 | -3.40 | 0.03 | 0.02 | 0.03 | 0.03 | 0.001 | 2 - Likely Not Pathogenic |
| P1806A | c.5416C>G | p.P1806A | -4.58 | -2.58 | 0.08 | 0.02 | 0.03 | 0.03 | 0.002 | 2 - Likely Not Pathogenic |
| V1808A | c.5423T>C | p.V1808A | -3.03 | -1.03 | 0.36 | 0.02 | 0.29 | 0.29 | 0.128 | NA - node 3 |
| V1809F | c.5425G>T | p.V1809F | 26.59 | 6.91 | 1000 | 0.02 | 0.03 | 0.03 | 0.969 | NA - node 3 |
| V1809A | c.5426T>C | p.V1809A | 1.43 | 0.00 | 1.00 | 0.02 | 0.29 | 0.29 | 0.290 | NA - node 3 |
| V1810G | c.5429T>G | p.V1810G | 8.72 | 6.72 | 832.6 | 0.02 | 0.81 | 0.81 | 1.000 | 4 - Likely Pathogenic |
| Q1811R | c.5432A>G | p.Q1811R | 14.66 | 6.91 | 1000 | 0.02 | 0.03 | 0.03 | 0.969 | NA - node 3 |
| P1812A | c.5434C>G | p.P1812A | -3.97 | -1.97 | 0.14 | 0.02 | 0.03 | 0.03 | 0.004 | 2 - Likely Not Pathogenic |
| D1813H | c.5437G>C | p.D1813H | -4.29 | -2.29 | 0.10 | 0.02 | 0.03 | 0.03 | 0.003 | 2 - Likely Not Pathogenic |
| D1818G | c.5453A>G | p.D1818G | -4.20 | -2.20 | 0.11 | 0.02 | 0.03 | 0.03 | 0.003 | 2 - Likely Not Pathogenic |
| N1819Y | c.5455A>T | p.N1819Y | -4.68 | -2.68 | 0.07 | 0.02 | 0.03 | 0.03 | 0.002 | 2 - Likely Not Pathogenic |
| N1819S | c.5456A>G | p.N1819S | -4.15 | -2.15 | 0.12 | 0.02 | 0.03 | 0.03 | 0.004 | 2 - Likely Not Pathogenic |
| A1823T | c.5467G>A | p.A1823T | -5.37 | -3.37 | 0.03 | 0.34 | 0.03 | 0.34 | 0.001 | NA - node 4 |
| Q1826H | c.5478G>C | p.Q1826H | -4.14 | -2.14 | 0.12 | 0.02 | 0.03 | 0.03 | 0.004 | 2 - Likely Not Pathogenic |
| A1830T | c.5488G>A | p.A1830T | -3.97 | -1.97 | 0.14 | 0.02 | 0.03 | 0.03 | 0.004 | 2 - Likely Not Pathogenic |
| V1833M | c.5497G>A | p.V1833M | 3.50 | 1.50 | 4.47 | 0.02 | 0.03 | 0.03 | 0.121 | NA - node 3 |
| R1835Q | c.5504G>A | p.R1835Q | -4.54 | -2.54 | 0.08 | 0.02 | 0.03 | 0.03 | 0.002 | 2 - Likely Not Pathogenic |
| R1835P | c.5504G>C | p.R1835P | -5.14 | -3.14 | 0.04 | 0.02 | 0.29 | 0.29 | 0.017 | NA - node 3 |
| E1836K | c.5506G>A | p.E1836K | 4.77 | 2.77 | 15.89 | 0.02 | 0.03 | 0.03 | 0.330 | NA - node 3 |
| W1837R | c.5509T>C | p.W1837R | 25.11 | 6.91 | 1000 | 0.02 | 0.81 | 0.81 | 1.000 | 4 - Likely Pathogenic |
| W1837G | c.5509T>G | p.W1837G | 5.95 | 3.95 | 51.88 | 0.02 | 0.81 | 0.81 | 0.995 | 4 - Likely Pathogenic |
| W1837C | c.5511G>C | p.W1837C | 5.15 | 3.15 | 23.24 | 0.02 | 0.81 | 0.81 | 0.990 | 4 - Likely Pathogenic |
| S1841R | c.5521A>C | p.S1841R | 5.78 | 3.78 | 43.98 | 0.02 | 0.81 | 0.81 | 0.995 | 4 - Likely Pathogenic |
| S1841N | c.5522G>A | p.S1841N | 13.84 | 6.91 | 1000 | 0.02 | 0.66 | 0.66 | 0.999 | 4 - Likely Pathogenic |
| A1843P | c.5527G>C | p.A1843P | 5.01 | 3.01 | 20.28 | 0.02 | 0.03 | 0.03 | 0.385 | NA - node 3 |
| L1844P | c.5531T>C | p.L1844P | -5.31 | -3.31 | 0.04 | 0.02 | 0.03 | 0.03 | 0.001 | 2 - Likely Not Pathogenic |
| D1851E | c.5553C>A | p.D1851E | -4.30 | -2.30 | 0.10 | 0.02 | 0.03 | 0.03 | 0.003 | 2 - Likely Not Pathogenic |
| Y1853C | c.5558A>G | p.Y1853C | 9.46 | 6.91 | 1000 | 0.02 | 0.81 | 0.81 | 1.000 | 4 - Likely Pathogenic |
| L1854P | c.5561T>C | p.L1854P | 7.66 | 5.66 | 286.7 | 0.02 | 0.29 | 0.29 | 0.992 | 4 - Likely Pathogenic |
| P1856S | c.5566C>T | p.P1856S | -4.59 | -2.59 | 0.07 | 0.02 | 0.03 | 0.03 | 0.002 | 2 - Likely Not Pathogenic |
| H1860P | c.5579A>C | p.H1860P | -5.20 | -3.20 | 0.04 | 0.02 | 0.03 | 0.03 | 0.001 | 2 - Likely Not Pathogenic |
| H1862L | c.5585A>T | p.H1862L | -4.30 | -2.30 | 0.10 | 0.02 | 0.03 | 0.03 | 0.003 | 2 - Likely Not Pathogenic |

NA – not applicable. Node 3 and 4 refer to the nodes in the decision tree of Figure 1, which explains why the sequence variant remains ‘unclassified’.

